# Adeno-Associated Virus-Induced Neurotoxicity is Prevented by CpG Depletion

**DOI:** 10.1101/2025.10.24.684017

**Authors:** Connor E. Dunn, Brigett V. Carvajal, Jie Ma, Guadalupe Castañeda-Hernandez, Ai Vy Nguyen, Caroline Jung, Lisa M. Boulanger

## Abstract

Adeno-associated viruses (AAVs) are the vector of choice for gene delivery to the nervous system. While AAVs have a strong safety profile, recent studies show that AAV causes dendritic loss and synaptic weakening in mouse somatosensory cortex, impacts that are prevented by systemic administration of blockers of Toll-like receptor 9 (TLR9), an innate immunoreceptor that detects unmethylated cytosine-guanine (CpG) motifs. However, TLR9 blockers are immunosuppressive and costly. To realize AAV’s full potential, it is critical to identify strategies to protect neurons without compromising immunity. Here we find that partially depleting CpG motifs from the AAV genome prevents AAV-induced dendritic loss and synaptic weakening. CpG depletion of transgene and intronic regions via codon-optimized synonymous mutations was protective and did not impair transgene expression *in vivo*. To facilitate the production and use of lower-CpG AAVs, we created a web-facing application, CpG-Assist Tool (CpG-AT), which allows researchers to quantify the CpG content of any sequence, explore the CpG content of commonly used AAV components, and generate CpG-depleted coding and non-coding sequences. This study identifies CpG depletion as a practical strategy to prevent AAV-induced neural circuit disruption, and provides tools to facilitate CpG reduction to enhance the safety and efficacy of AAV-mediated gene delivery.

## Introduction

Adeno-associated viruses (AAVs) are non-enveloped, single stranded DNA viruses that are widely used for gene delivery. AAVs are non-pathogenic and cannot replicate in the absence of a helper virus^1^, contributing to their strong safety profile. The AAV genome can persist as an extrachromosomal episome in the nucleus of transduced cells, driving long-term transgene expression^2,3^. Recombinant AAVs have been modified to replace viral genes with a variety of transgenes, as well as promoters, enhancers, and other *cis* elements to strengthen and target transgene expression^4,5^. Recombinant AAVs can also be engineered to have either broad or specific tropism, making them powerful and flexible tools for a wide variety of research and clinical applications^6^.

In most tissues, the host immune response can pose significant challenges to AAV-mediated gene delivery. Peripheral immune responses against diverse elements of AAV preparations have been observed, including the AAV genome^7,8^, capsid^9^, expressed transgene^10^, contaminants^11^, and process- and production-related impurities^12^. Immune responses to AAV-mediated gene delivery can include neutralizing antibodies, complement activation, and recruitment and/or activation of CD8^+^ and CD4^+^ T cells, B cells, NK cells, and dendritic cells^9,13–15^. As a result of the immune response, viral genomes can be cleared^16^, transgene expression can be curtailed or eliminated^17–19^, and tissue damage can occur^14,20–22^. Because of the significant impacts of peripheral immunity, patients may be excluded from AAV-based clinical trials or treatments if they have pre-existing anti-AAV antibodies^23–25^, and repeated dosing may not be effective^26^. Recognition of unmethylated cytosine-guanine (CpG) motifs in the AAV genome by the innate immunoreceptor Toll-like receptor 9 (TLR9) in particular plays a central role in initiating both innate and adaptive immune responses after AAV-mediated gene delivery^7,8,27,28^. Consistent with TLR9’s role in initiating transgene-limiting immune responses, reducing TLR9 activation is associated with more robust and persistent transgene expression^18,29^.

In contrast, administration of AAV in the central nervous system (CNS) is not associated with the same robust inflammatory response that is seen in the periphery. The immune privileged status of the CNS dampens transgene-limiting inflammation and allows more stable and prolonged transgene expression in nervous system cells^30–32^. As a result, AAV is widely used in neuroscience research, and is currently the basis of multiple FDA-approved CNS gene therapies, including Luxturna, a treatment for *RPE65* mutation-associated retinal dystrophy, Zolgensma, a treatment for spinal motor atrophy, and Kebilidi, a treatment for aromatic L-amino acid decarboxylase deficiency^33–35^. AAV is also the vector in many ongoing clinical trials to treat diverse CNS disorders.

While AAV does not cause significant inflammation in the CNS, recent studies have identified other unwanted impacts of AAV-mediated gene delivery in the CNS in animal models. AAV administration via the cerebral spinal fluid or via high dose intravenous injection is associated with neurodegeneration in the dorsal root ganglia in non-human primates and pigs^36–38^. Intracranial administration of AAV is associated with neurotoxicity in non-human primates^39^, and disrupts the blood-brain barrier and is associated with neuronal death in adult mice^21^. Subretinal administration of AAV causes retinal pigment epithelium toxicity in neonatal mice^20^. AAV-mediated gene delivery in adult mouse hippocampus is associated with death of neural progenitor cells and immature dentate granule cells, and impaired neurogenesis^40^. Isolated inverted terminal repeats (ITRs) from the same AAVs are also sufficient to cause death of neural progenitor cells *in vitro*^40^, suggesting that an element of AAV’s DNA genome can initiate neuronal damage.

One potential trigger for AAV-induced neuronal damage is unmethylated CpG DNA. We recently found that AAV-mediated gene delivery reduces dendritic complexity and disrupts synaptic transmission in adult mouse somatosensory cortex. These changes are not restricted to a particular AAV serotype, promoter, or encoded transgene, suggesting they are caused by a common feature of diverse AAVs^41^. One such feature, unmethylated CpG DNA, is found in essentially all AAV genomes^42–44^ and is detected by TLR9. Strikingly, both structural and functional impacts of AAV in the brain are prevented by systemic administration of ODN 2088, a synthetic TLR9 antagonist^41^. The efficacy of TLR9 inhibitors in preventing AAV-induced dendrite loss and synaptic weakening suggests that CpG dinucleotides in the AAV genome may activate TLR9 to drive circuit damage.

To realize the full potential of AAV-based research tools and therapies, it is critical to identify strategies to mitigate the unwanted effects of AAV on neurons. While systemic treatment with a TLR9 inhibitor can protect cortical neurons^41^, this approach is broadly immunosuppressive, and is not cost-effective, particularly in larger animals. A complementary approach to reduce or prevent host TLR9 responses is to remove TLR9 ligands from the AAV genome. In the periphery, depleting CpG dinucleotides from AAV DNA can combat some, but not all, immune responses to AAV-mediated gene delivery^18,19,29,45^. We hypothesized that CpG depletion might reduce AAV-induced circuit damage in the mammalian CNS.

To test this hypothesis, we reduced the CpG content of transgene and intronic regions of the AAV genome. We found that CpG depletion completely prevents AAV-induced loss of dendritic complexity and disruption of synaptic transmission, without compromising transgene expression. We also found that AAV genomic elements that are commonly used in the CNS vary widely in their CpG content, suggesting that some may be more likely to trigger TLR9-mediated neuronal damage than others. To help AAV users minimize the CpG content of their vectors, we created a web-facing, freely available application, CpG-Assist Tool (CpG-AT). CpG-AT allows users to browse the CpG content of a large library of AAV elements and quantify or deplete the CpG content of user-supplied DNA sequences. Since we find that CpG depletion is an effective strategy to protect neurons during AAV-mediated gene delivery, CpG-AT could facilitate the design and dissemination of safer and more effective AAV-based gene delivery systems.

## Results

### Depletion of CpG motifs from AAV1-*pCAG*-*FLEX*-*EGFP*

To determine if CpG depletion is a viable strategy to reduce AAV-induced circuit disruption in the CNS, we removed a subset of CpG dinucleotides from the genome of AAV1-*pCAG*-*FLEX*-*EGFP* (**Figure 1**). This virus caused both dendritic loss and synaptic weakening in mouse somatosensory cortex in our previous study^41^. We depleted CpGs from the transgene (*EGFP*) and intronic regions, including the chimeric intron (**Figure 1A-C**; see Materials and Methods for details). We did not modify the inverted terminal repeats (ITRs), CAG promoter, WPRE, or loxP regions, since modifying these regulatory regions may negatively impact transgene expression^46^.

**Figure 1.**
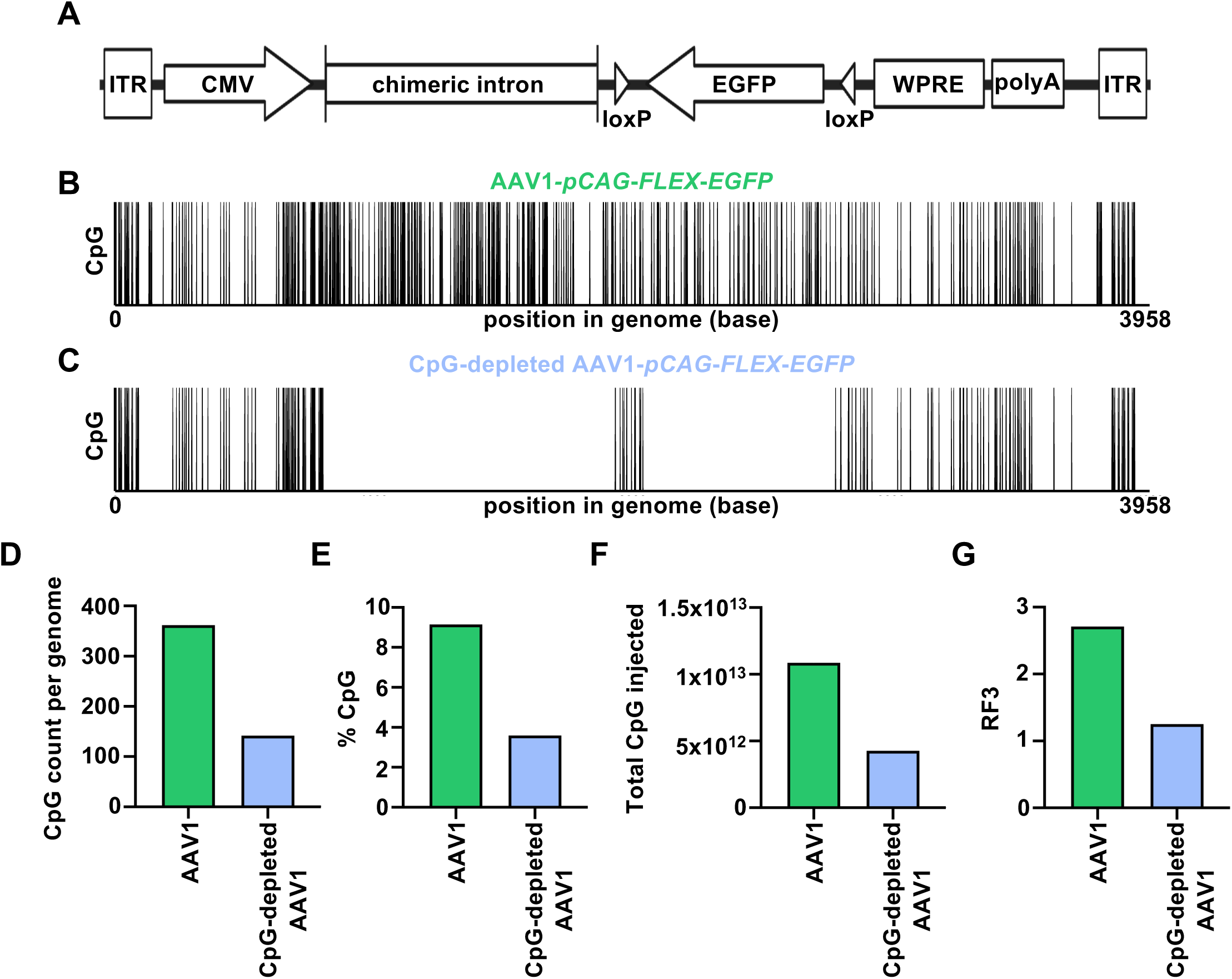
Depleting CpG dinucleotide motifs from coding and intronic regions of AAV1-*pCAG*-*FLEX*-*EGFP*. **(A)** Domain structure of AAV1-*pCAG*-*FLEX*-*EGFP.* **(B)** CpG motifs in the AAV1-*pCAG*-*FLEX EGFP* and **(C)** CpG-depleted AAV1-*pCAG*-*FLEX*-*EGFP* genomes. Each vertical line represents a single CpG dinucleotide. CpG dinucleotides were removed from transgene coding and intronic regions. **(D-G)** Measures of CpG content **(D-F)** and predicted TLR9 activating potential^50^ **(G)** before (green) and after (blue) CpG depletion.

A central goal when depleting CpGs is to minimize effects on transgene structure, function, and expression. To this end, we replaced CpG dinucleotide-containing codons (e.g., CGG) and CpG dinucleotide-spanning codons (e.g., CUC-GGG) in the transgene coding region by creating synonymous, species-specific, codon-optimized mutations. To determine which codon(s) to use, we consulted updated codon usage tables found in the High-performance Integrated Virtual Environment-Codon Usage Tables (HIVE-CUTs) database^47^. We replaced CpG-containing and CpG-spanning dinucleotides with codons that did not change the encoded amino acids (synonymous mutations) and had the highest usage in mice. If the highest usage codon was being replaced, we replaced it with the next-highest usage codon. In intronic regions, we removed CpG dinucleotides by replacing the C with an A. We have made a modified version of this CpG depletion pipeline for both coding and intronic regions available via our online application (see CpG-Assist Tool, below).

Applying this CpG depletion pipeline decreased the total number of CpG dinucleotides from 362 in AAV1-*pCAG*-*FLEX*-*EGFP* (“unmodified AAV”) to 142 in the CpG-depleted AAV1*-pCAG-FLEX-EGFP* (“CpG-depleted AAV”), a ∼60% reduction (**Figure 1D**). Because the length of the AAV genome was unaltered, the effect on the percent CpG content was also a ∼60% reduction: unmodified AAV contained 9.15% CpG dinucleotides, whereas CpG-depleted AAV contained 3.59% CpG dinucleotides (**Figure 1E**). Because the same amount of each virus was injected (1×10^10^ vg), the total number of CpGs injected per animal also dropped proportionally (**Figure 1F**).

TLR9 recognizes unmethylated CpG dinucleotides in the context of their flanking nucleotides, which influence if TLR9 will be activated or inhibited by a given CpG motif^48,49^. Therefore, to most accurately estimate the TLR9-activating potential of a given AAV genome, it is important to consider not only the number or percent of CpGs, but also their nucleotide context. To that end, we used a metric, termed RF3, created by Wright^50^, that classifies specific CpGs as stimulatory or inhibitory for TLR9, and also considers the total number of CpG motifs and overall sequence length. The RF3 value of the CpG-depleted AAV (RF3 = 1.25) is less than half that of unmodified AAV (RF3 = 2.71) (**Figure 1G**). Thus removing CpGs in the intronic and coding regions alone dramatically reduced the number and percentage of CpGs as well as the predicted TLR9 activating potential of the AAV genome, as estimated by RF3.

### CpG-depleted AAV1-*pCAG*-*FLEX*-*EGFP* drives robust transgene expression

While the substitutions made to replace CpG motifs are conservative, meaning they do not change the encoded amino acids, they could still affect transgene expression^51^. Intronic mutations can also affect transgene expression^52^. To determine if CpG depletion disrupts transgene expression, we quantified EGFP levels in the brains of adult mice that received a unilateral intraparenchymal injection of 1×10^10^ viral genomes (vg) of either unmodified or CpG-depleted AAV into primary somatosensory cortex (S1; see Materials and Methods). The LoxP sites in the *FLEX* plasmid require CRE recombinase for efficient recombination^53^. Therefore, to induce transgene expression, we injected unmodified or CpG-depleted AAV into *Slc17a7-IRES2-Cre-D* mice, which express Cre recombinase under the control of the *Slc17a7* promoter. This promoter drives transgene expression in glutamatergic neurons in cortex, including S1^54^. Twenty-one days after AAV injection (21 dpi), levels of EGFP protein were quantified in lysates of S1 via Western blotting, using GAPDH as a loading control. EGFP levels in injected hemispheres of cortex were indistinguishable in mice receiving either unmodified or CpG-depleted AAV (**Figure 2A-B**). These results suggest that CpG depletion of the AAV1-*pCAG*-*FLEX*-*EGFP* genome did not disrupt transgene expression in the CNS *in vivo*.

**Figure 2.**
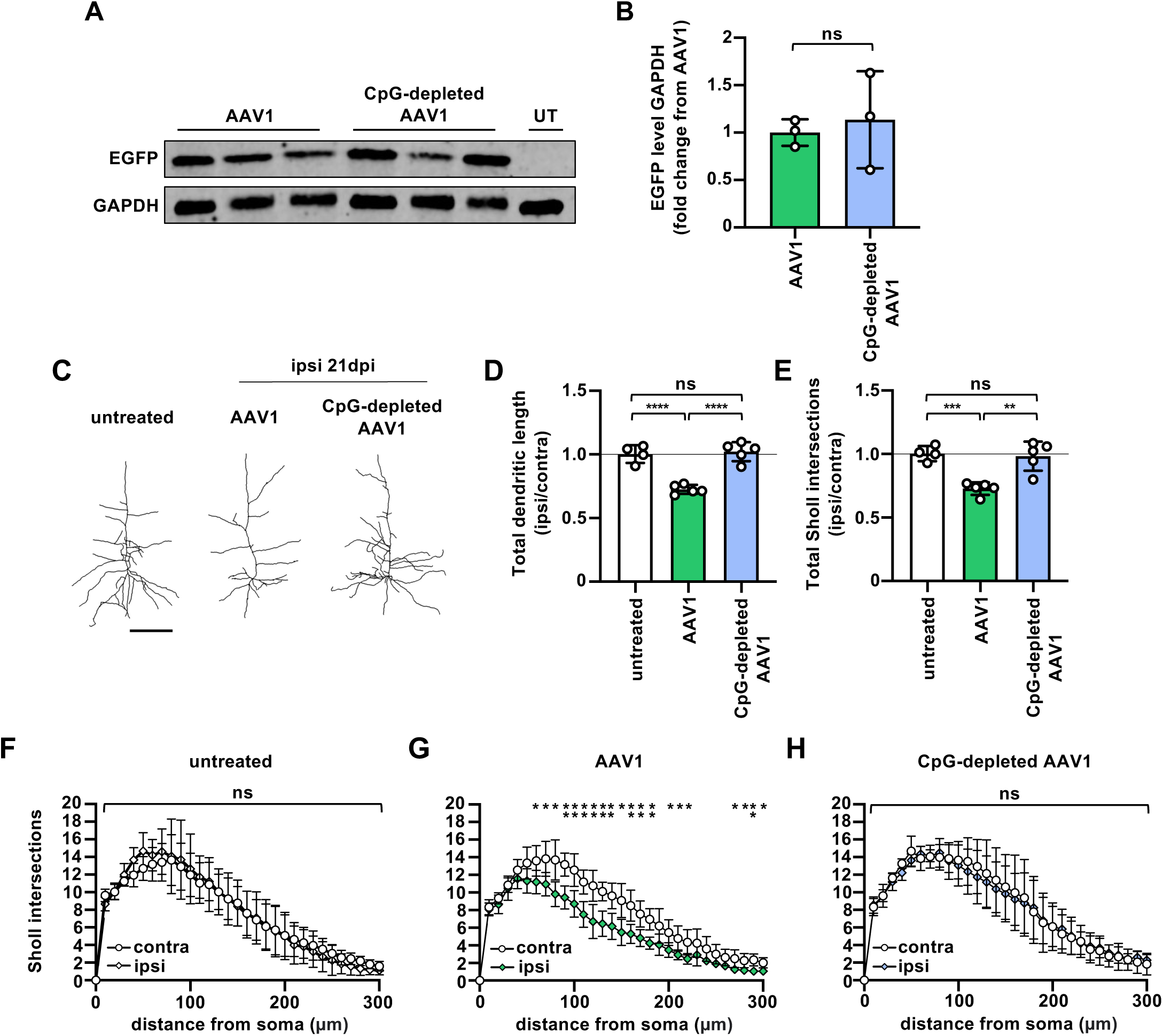
CpG depletion does not alter transgene expression but prevents AAV-induced dendritic loss. **(A)** Immunoblot of S1 homogenates probed with antibodies against transgene product EGFP and loading control GAPDH following injection of 1×10^10^ vg of unmodified AAV1-pCAG-FLEX-EGFP or CpG-depleted AAV1-pCAG-FLEX-EGFP into Slc17a7-IRES2-Cre-D mice. Each lane contains samples from a different animal. UT, untreated. **(B)** Densitometry shows that EGFP protein levels are indistinguishable in ipsilateral S1 of Slc17a7-IRES2-Cre-D mice injected with either CpG-depleted or unmodified virus. **(C)** Representative cortical neurons from untreated wild type mice or ipsilateral (injected) hemispheres of wild type mice injected with 1×10^10^ vg of either unmodified or CpG-depleted AAV1-pCAG-FLEX-EGFP. Scale bar, 100 μm. **(D)** Total dendritic length and **(E)** total Sholl intersections of cortical neurons from untreated wild type mice and wild type mice injected with 1×10^10^ vg of either unmodified or CpG-depleted AAV1-pCAG-FLEX-EGFP. Values represented as a ratio of injected (ipsi) normalized to uninjected (contra) hemispheres. **(F-H)** Sholl intersections plotted as a function of distance from the soma for cortical neurons from the same animals shown in **(E)**. **(D-E)**, n=4 untreated control mice, 10-14 neurons per animal (average = 12 cells); n=5 mice injected with unmodified AAV1-pCAG-FLEX-EGFP, 10-14 neurons per animal (average = 12 cells); n=5 for mice injected with CpG-depleted AAV1-pCAG-FLEX-EGFP, 12-15 neurons per animal (average = 13 cells). Data in (**B-E)** represented as mean ± SD. \*\*\*\**p* < .0001; \*\*\**p* < .001; \*\**p* < .01; ns, not significant. In **(G)**, single * indicate p<0.05, vertically stacked * indicate p<0.01. Statistical analysis: in **(B, F-H)**, two-tailed unpaired t-test; in **(D-E)**, one-way ANOVA with Tukey’s multiple comparison test.

### CpG depletion of AAV1-*pCAG*-*FLEX*-*EGFP* prevents loss of cortical dendritic complexity

Intraparenchymal injection of unmodified AAV1-*pCAG*-*FLEX*-*EGFP* into S1 of WT mice reduces dendritic length and complexity in cortical pyramidal neurons. These neuroanatomical changes are prevented by systemic administration of the TLR9 antagonist ODN 2088^41^, suggesting they may be mediated by TLR9. To determine if depletion of TLR9 ligands (CpGs) from the AAV genome can reduce AAV-induced dendritic loss, we injected either unmodified or CpG-depleted AAV (at the same dose, 1×10^10^ vg) into S1 of 8 to 10-week-old wild type C57BL/6J mice. WT mice were used to avoid any changes that might be induced by transgene expression, since transgene expression is not required for AAV-induced dendritic loss^41^. We quantified dendritic length and complexity of individual layer IV-VI pyramidal cortical neurons at 21 dpi, using a DiOlistic labeling approach^55^. Tungsten bullets were coated in a lipophilic dye (DiO) and delivered to fixed tissue sections using a gene gun at an appropriate distance and pressure to sparsely label individual neurons (see Materials and Methods; **Supplementary Figure 1A-B**)^55^. Total dendritic length was measured, and Sholl analysis^56^ was used to quantify dendritic complexity.

In uninjected contralateral hemispheres, dendritic length and Sholl intersections were indistinguishable from values in age-matched untreated controls **(Supplementary Figure 1C-E)**, suggesting that any effects of AAV on these metrics are restricted to the injected hemisphere, consistent with previous results^41^. Therefore in subsequent analyses, total dendritic length and total Sholl intersection values are represented as ipsilateral/contralateral ratios, to control for any inter-animal variability. Unmodified AAV caused a significant decrease in dendritic length and complexity in the injected ipsilateral hemisphere (**Figure 2D-H**), consistent with previous results using the same virus, dose, age range, and strain of mice^41^. In striking contrast, in animals injected with CpG-depleted AAV, both total dendritic length (**Figure 2D**) and Sholl intersections (**Figure 2E-H**) in the injected hemisphere are significantly increased compared to mice injected with the unmodified AAV and are indistinguishable from untreated control animals (**Figure 2D-E**; for distributions of individual cells see **Supplementary Figure 1F-G)**. In animals injected with unmodified AAV, loss of Sholl intersections is distributed throughout the dendritic arbor (**Figure 2G**), and in animals injected with CpG-depleted AAV, dendrites are protected at all distances from the soma (**Figure 2H**). Taken together, these results suggest that partial CpG depletion fully prevents the loss of dendritic length and complexity caused by AAV1*-pCAG-FLEX-EGFP*.

### CpG depletion of AAV1-*pCAG*-*FLEX*-*EGFP* prevents disruption of synaptic transmission

We previously found that injection of 1×10^10^ vg of AAV1-*pCAG*-*FLEX*-*EGFP* into mouse S1 disrupts synaptic transmission, apparent as a reduction in the frequency of miniature excitatory postsynaptic currents (mEPSCs)^41^. mEPSCs represent responses to spontaneous release of synaptic vesicles at individual synapses, and a decrease in mEPSC frequency is consistent with a decrease in either the presynaptic probability of release or the number of functional synapses. In the current study, we find that CpG depletion of the AAV genome (**Figure 1**) can prevent AAV-induced dendritic loss (**Figure 2**). To determine if CpG depletion can also protect against functional synaptic impacts of AAV, we injected 1×10^10^ vg of unmodified or CpG-depleted AAV1*-pCAG-FLEX-EGFP* into S1 of 8 to 10-week-old wild type C57BL/6J mice. At 21 dpi, whole cell patch-clamp recordings were performed, and mEPSCs recorded (**Figure 3A)**. In animals injected with unmodified AAV, mEPSC frequency was reduced relative to uninjected controls, consistent with previous results^41^. In animals injected with CpG-depleted AAV, in contrast, mEPSC frequency was significantly increased relative to animals injected with unmodified AAV and was indistinguishable from untreated controls (**Figure 3B-C)**. Thus depleting CpG motifs can prevent both structural and functional impacts of AAV-mediated gene delivery in the CNS.

**Figure 3.**
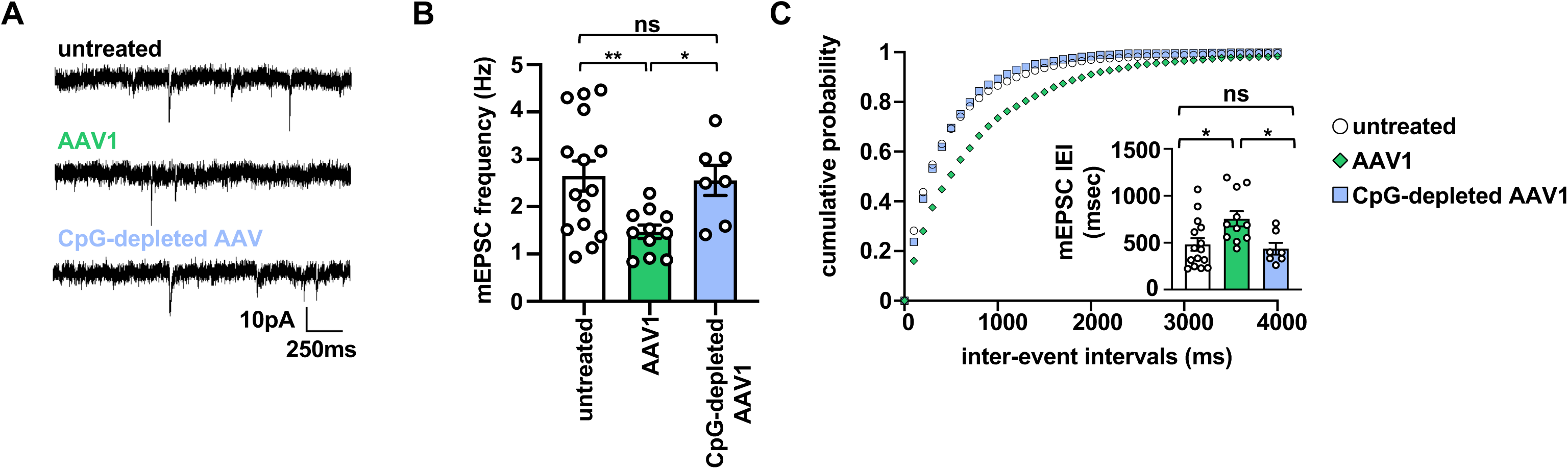
CpG depletion prevents AAV-induced changes in synaptic transmission. **(A)** Example whole cell recordings from S1 pyramidal neurons in acute brain slices from an untreated control animal or from the ipsilateral hemispheres of animals injected with 1×10^10^ vg of either unmodified or CpG-depleted AAV1-*pCAG*-*FLEX*-*EGFP*. **(B-C)** Administration of unmodified AAV reduces mEPSC frequency and increases the inter-event interval (IEI). CpG depletion significantly increases mEPSC frequency and decreases IEI relative to animals injected with unmodified AAV to levels that are indistinguishable from untreated controls. Data in **(B-C)** represented as mean ± SD. \*\**p* < .01; \**p* < .05; ns, not significant. Statistical analysis in **(B)**, Brown-Forsythe one-way ANOVA test with Dunnett’s T3 multiple comparison test; **(C)**, one-way ANOVA with Tukey’s multiple comparison test.

### CpG content of commonly used AAV components varies widely

The above results suggest that partially depleting CpGs from the AAV genome can prevent AAV’s detrimental impacts on the structure and function of cortical circuitry. Directed CpG depletion requires modification of plasmids prior to viral synthesis, a process that can make virus production more time-consuming and costly. To determine if there are existing, turnkey AAV genome components that are relatively low in CpG content, we assembled a library of commonly used DNA sequences, with an emphasis on elements used in the CNS (for sources of DNA sequences, see **Supplementary Table 1**). This library includes 34 promoters/enhancers (12 promoters with broad tropism, 20 tissue-specific promoters, and 2 inducible promoters); 90 transgenes (31 fluorescent tags/tracers, 17 optogenetic transgenes, 4 chemogenetic transgenes, 32 biosensors, 3 recombinases, and 3 other); 16 miscellaneous components (6 regulatory elements, 5 tags, 4 linkers, and 1 histone); and ITRs from 8 different AAV serotypes. Although this library is not exhaustive, it includes many AAV components that are commonly used in the CNS. This library, along with the corresponding quantification of CpG content, is available via a web-facing application (see CpG-Assist Tool application, below).

We quantified the CpG content of these sequences using three metrics: total CpG dinucleotides, % CpG dinucleotides, and RF3, a metric intended to estimate predicted TLR9 activation by differently weighting stimulatory and inhibitory CpG motifs, while also considering the total number of CpGs and the total length of the sequence^50^. The results of these analyses are provided in **Supplementary Table 1** and are shown in **Figure 4** (RF3 for transgenes, promoters, enhancers, and miscellaneous components), **Supplementary Figure 2** (% CpG dinucleotides for transgenes, promoters, enhancers, and miscellaneous components), **Supplementary Figure 3** (total CpG dinucleotides for transgenes, promoters, enhancers, and miscellaneous components), and **Supplementary Figure 4** (RF3, total CpG dinucleotides and % CpG for ITRs. The CpG content of ITRs may not be as relevant to vector design, as the AAV2 ITR is almost uniformly used in recombinant AAV vectors, likely due to its strong transcriptional/promoter activity^57^). We found there is remarkable variation in CpG content, even among components that have a related function, such as broad promoters/enhancers or fluorescent tracers. Thus, it is possible to select or design an AAV that performs a given function, such as expressing a fluorescent tag under the control of a broad promoter, using components with relatively low CpG content. This approach has the benefit that it achieves lower CpG content without requiring modification of the DNA via targeted CpG depletion, and therefore may be faster, more cost-effective, and avoid the potential for changes in transgene levels or localization as a consequence of introducing synonymous mutations.

**Figure 4.**
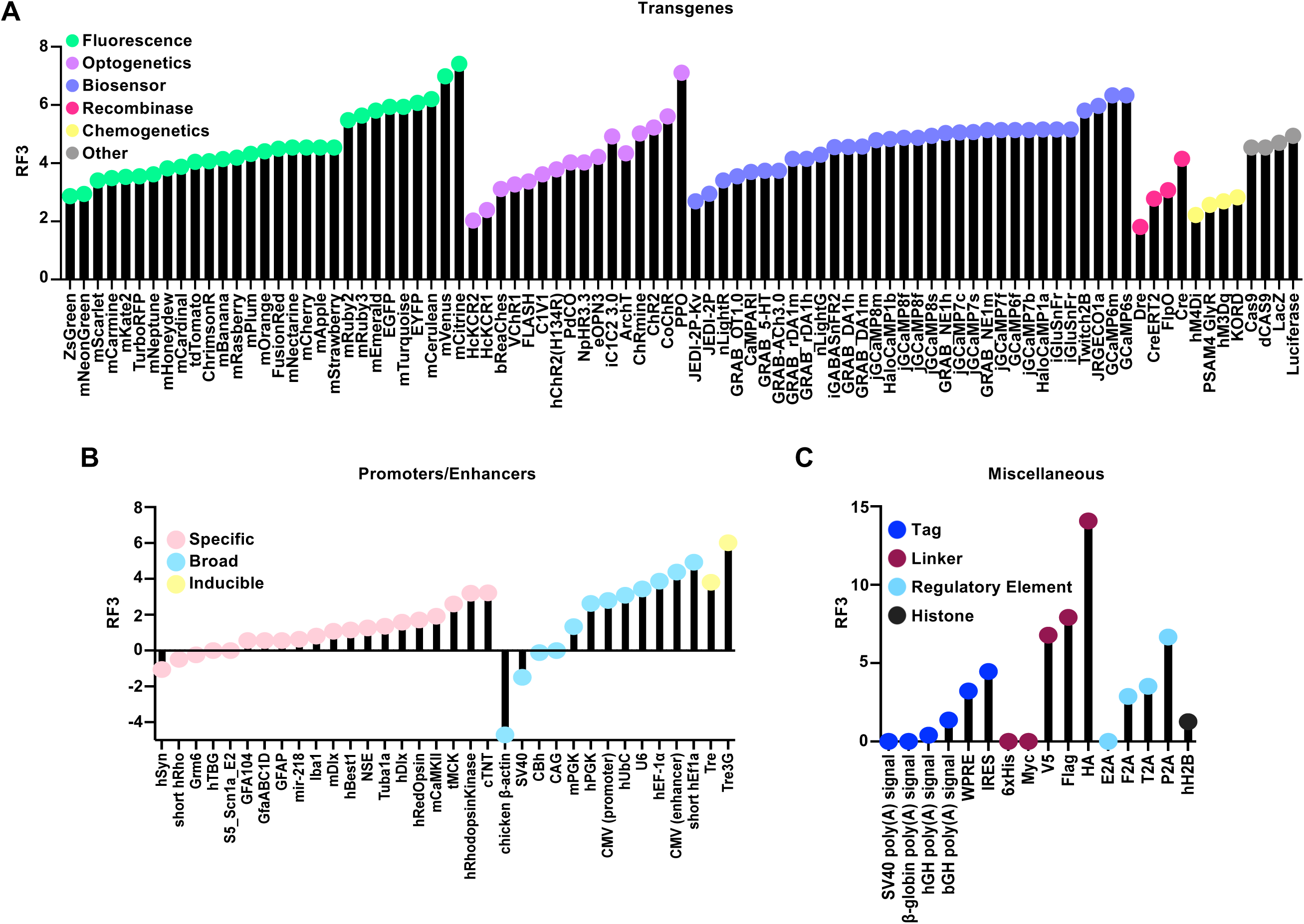
RF3 values for commonly used AAV transgenes, promoters, enhancers, and miscellaneous components. RF3^50^ values for **(A)** transgenes classified by general function: fluorescence, optogenetics, biosensor, recombinase, chemogenetics, and other; **(B)** promoters/enhancers classified by broad, specific, and inducible; and **(C)** miscellaneous components classified by tags, linkers, regulatory elements, and histones.

### CpG-Assist Tool: a web-facing, interactive application to quantify, reduce, and explore the CpG content of existing and user-supplied DNA sequences

With the goal of helping researchers combat AAV-induced changes caused by activation of TLR9 in the CNS and beyond, we developed a publicly accessible, web-based application (https://cpgat.princeton.edu/). There are three primary functions of this application, which we named CpG-Assist Tool (CpG-AT).

The first function of CpG-AT is to quantify the CpG content of user-submitted DNA sequences. This tool will quantify the total CpG dinucleotides, % CpG dinucleotides, and RF3 value of any single-stranded DNA sequence. Users can visualize the CpG content of their sequences using a barcode plot, as in **Figure 1B-C**, which displays CpG dinucleotides across the length of the sequence, making it easy to identify regions of relatively high CpG content that may be candidates for replacement or CpG depletion. The second function of CpG-AT is to allow users to browse a library of 148 commonly used AAV components alongside their quantified CpG content. Users can explore how the CpG content of different AAV components varies, or search for a specific component of interest.

The third function of CpG-AT is to provide a template to deplete CpG motifs from a given sequence using a version of the pipeline we developed for this study. Sequences can either be intronic or coding. For coding sequences, users can specify the host species, and the CpG depletion tool will consult codon usage patterns of that species^47^ to create codon-optimized synonymous mutations. The current version of CpG-AT supports codon optimization for mice, rats, and humans. The CpG depletion pipeline outputs a new, CpG-depleted sequence, a side-by-side visualization of the CpG content of the unmodified and CpG-depleted sequences, and if a coding sequence is used, the amino acid sequences of the unmodified and CpG-depleted versions, to verify there are no changes at the amino acid level. Intronic sequences are CpG-depleted by changing the C in CG motifs to a G, if the preceding base is not a C. In cases where the preceding base is a C, the C from the CG motif is changed to an A. Substituting Gs for Cs where possible during CpG depletion has the advantage that it keeps the GC content of CpG-depleted intronic regions as consistent as possible with original sequences, which may help with virus production and/or transgene expression^51,58^. Overall, the CpG-AT application allows quantification, visualization and depletion of putative TLR9 ligands in the AAV genome. Our *in vivo* validation demonstrates the protective value of CpG depletion for AAV use in the nervous system, and therefore the CpG-AT application can facilitate the selection and design of AAVs that are less likely to elicit damaging TLR9 responses.

## Discussion

AAV is currently the vector of choice for viral gene delivery and has an overall strong safety profile. However, AAV-mediated gene delivery can reduce dendritic complexity and weaken synaptic transmission in mouse somatosensory cortex^41^. Here we show that depleting TLR9 ligands (CpG motifs) from the AAV genome prevents these AAV-induced changes in circuit structure and function. The protection afforded by CpG depletion is not associated with a drop in transgene expression in the CNS. The effectiveness of removing TLR9 ligands seen here, as well as the efficacy of systemic TLR9 inhibitors in our previous study^41^, together strongly suggest that AAV causes dendritic loss and synaptic weakening *in vivo* via TLR9-mediated detection of CpG motifs in the AAV genome. The current study identifies CpG depletion as a practical approach to improve the safety of AAV as a gene delivery vector in the CNS.

In addition to CpG depletion, other strategies may dampen or avert harmful responses to AAV-mediated gene delivery in the CNS. Injecting fewer AAV genomes can be protective, though lowering the dose sufficiently to be protective may compromise transgene expression^21,41,59^. High viral titers may be more safely delivered to the CNS by co-administrating TLR9 inhibitors (like ODN 2088)^41,60^, though there are three issues with this approach. First, systemic administration of TLR9 inhibitors is immunosuppressive, which may be undesirable in experimental and clinical settings. Second, systemic administration of ODN 2088 may not be cost-effective, particularly in larger animals. Third, TLR9 signaling may have functional roles that extend beyond inflammation, particularly in the nervous system, and these non-inflammatory roles of TLR9 may be affected by drugs that target TLR9. For example, one recent study found that neuron-specific deletion of TLR9 inhibits the formation of perineuronal nets in the hippocampus and impairs contextual fear conditioning. TLR9 was also essential for maintaining neuronal genomic stability in the face of ongoing neural activity^61^. Thus TLR9 inhibition may have undesirable impacts beyond immunosuppression, particularly in the nervous system. Depleting CpGs from the AAV genome is a cost-effective neuroprotective approach that avoids immune and non-immune effects of TLR9 inhibition, while allowing injection of sufficient viral titers to support robust transgene expression.

This study is, to our knowledge, the first to test the efficacy of CpG depletion of the AAV genome as a protective approach in the CNS. Outside the CNS, CpG depletion has been used to combat TLR9-induced peripheral immune responses that can blunt transgene expression and cause tissue damage. For example, CpG depletion of the transgene region of the AAVrh32.33 vector significantly reduced transgene-specific CD8^+^ T cells and expression of CD4^+^ and MHCII in transduced skeletal muscle^18^. Depletion of CpG motifs from an AAV encoding coagulation factor IX (FIX) significantly reduced CD8^+^ T cell responses against the transgene when injected into a mouse model of hemophilia B^29^. In a recent phase 1/2 clinical trial, an AAV-based therapy encoding FIX was administered to patients with hemophilia B^19^. When the AAV was codon-optimized, CpG motifs were incidentally introduced into the vector, changes that correlated with a rapid loss of transgene expression. To test if CpG enrichment could be causing an immune response against the AAV vector, the FIX transgene was CpG-depleted and administered to mice, resulting in significantly lower levels of anti-AAV8 neutralizing antibodies compared to the unmodified vector^19^. Taken together, these and other studies suggest that CpG depletion can help circumvent AAV-induced immune responses and subsequent loss of transgene expression in the periphery. The beneficial effects of CpG depletion in our CNS study, and in the periphery, suggest that the CpG-AT application developed here may be broadly useful for the design of safe and effective AAVs for central as well as peripheral uses.

Here we show that CpG depletion can prevent AAV-induced dendrite loss and synaptic weakening in the CNS. Although AAV does not cause widespread cell death in the CNS, several recent studies have identified focal AAV-associated cell loss in the CNS. These include retinal pigment epithelium toxicity in neonatal mouse retina^20^, neural progenitor cell death and impaired neurogenesis in adult mouse hippocampus^40^, disruptions to the blood-brain barrier and neuronal death in adult mouse cortex^21^, neurotoxicity in the adult non-human primate thalamus^39^, and neurodegeneration in non-human primate and pig dorsal root ganglia^36–38^. Although some of these neurotoxic effects of AAV are thought to reflect non-immune mechanisms, including transgene toxicity^39,62^, for others the mechanisms remain unclear. Of note, the effects of AAV on hippocampal neurogenesis are mimicked by injecting only AAV’s inverted terminal repeats^40^, suggesting that suppression of neurogenesis is a response to AAV DNA. CpG DNA is cytotoxic to neurons in organotypic slice cultures *in vitro*^63^, supporting a model in which responses to CpGs in AAV’s DNA genome can be neurotoxic. It will be important to determine if TLR9 mediates these or other AAV-associated effects on the nervous system, and if CpG depletion would be beneficial.

The design of safer AAVs that minimize TLR9-mediated responses requires a metric that robustly and quantitatively predicts TLR9 activation by a given DNA genome. It remains an open question what metric (e.g., total CpG count, % CpG, RF3) is the most reliable predictor of TLR9 activation. There are cases in which a DNA sequence is low in one metric, and high in another. Of these metrics, only RF3 takes into account the fact that some CpGs are TLR9 agonists while others are TLR9 antagonists^50^. RF3 is based on six known TLR9 agonist and antagonist motifs; other CpGs are considered but assigned less weight. There are likely far more TLR9 agonist and antagonist sequences, since TLR9 ligand space remains incompletely mapped.

More work is needed to identify and properly weight a larger pool of TLR9 agonist and antagonist motifs, based on quantitative readouts of TLR9 activation, and incorporate these findings into the RF3 calculation. Commercially available TLR9 reporter cell lines could potentially be used to validate RF3 predictions and test novel oligodeoxynucleotide sequences^64^. Although the precise weighting of stimulatory and inhibitory motifs in calculating RF3 is preliminary, the sign and relative magnitude of RF3 values are consistent with empirical data. For example, viral and bacterial DNA more robustly activate human TLR9 than human DNA^50,65^, and on average have higher RF3 values (∼4.7 for the entire *E. coli* genome, ∼ 5 for the entire wildtype AAV2 genome) than the human genome (∼0.2 for the entire genome, ∼0.5 for a relatively CpG-rich region of chromosome 1)^50^. Increases in RF3 also correlated with blunted transgene expression and more immune-related complications in a clinical trial using AAVs carrying an optimized coagulation FIX^19,50,66–70^. Thus RF3, while preliminary, is likely the most accurate tool currently available to estimate TLR9-mediated responses to DNA sequences in mammalian cells.

In the current study, CpG depleting only the transgene coding and intronic regions was sufficient to fully prevent dendritic and synaptic changes. We did not deplete CpGs from the ITRs or promoter, and the genome overall retained ∼40% of its CpG content after depletion. The qualitative effects of this quantitative change in CpG content suggest that there is a threshold level of CpG content that induces TLR9-dependent changes in neurons. Such a thresholding system for TLR9 activation could help ensure that low levels of unmethylated host DNA are insufficient to trigger a harmful autoimmune reaction. Wright proposed an RF3 threshold of ∼1.34 for clinically relevant TLR9-dependent immune responses in the periphery. This threshold was inferred from transgene longevity and immune-related complications of 4 AAV-FIX vectors in the periphery in human subjects^19,66–70^. While the current study is in the CNS, and in mice, our unmodified AAV exceeded this proposed threshold (RF3 of 2.71), while our CpG-depleted AAV fell below the threshold (RF3 of 1.25). Further refinements to the RF3 calculation will be necessary to more precisely predict TLR9 responses, but an RF3 of 1.34 or lower may be a useful provisional target when designing AAVs.

While CpG depletion completely protected against dendritic impacts of AAV without altering transgene levels in this study, refinements to the CpG depletion process might further improve outcomes in other settings. First, to create our CpG-depleted virus, we converted C residues in CpG motifs in the promoter and coding region to As. The CpG-AT application instead allows users to convert these C residues to Gs when possible, thereby retaining the original GC content of the sequence, which may affect virus production and/or transgene expression levels^51,58^. Second, in our virus, all CpGs in the intronic and transgene regions were eliminated. However, some CpG motifs are TLR9 inhibitors, while others are TLR9 agonists, and a third category have effects on TLR9 that have not been quantified. It will be important to test if eliminating only the activating and uncategorized CpG motifs, but leaving the inhibitory motifs intact, provides the same or even additional protection. Alternatively, CpGs could be converted to inhibitory motifs, a change that would preserve the CG content of the sequence while more dramatically reducing the RF3 value of the AAV genome. It is likely that not all CpGs in the transgene region can be changed to inhibitory while maintaining synonymous mutations; in this case, a synonymous mutation removing the CpG can be used. A related technique, encoding longer, exogenous TLR9 inhibitors in the AAV genome, has been tested in animal models and can reduce innate immune activation and improve transgene expression^71,72^. Converting endogenous CpGs to inhibitory motifs has the advantage that it does not take up additional space in AAV’s limited genome capacity. *In silico* analysis shows that converting activating and uncategorized CpGs in the transgene and intronic regions of AAV1-*pCAG*-*FLEX*-*EGFP* to inhibitory CpGs gives this AAV an RF3 value of -2.59, a marked decrease from the RF3 of the unmodified virus (2.71) as well as our CpG-depleted virus (1.25).

In the periphery, activation of TLR9 results in parallel activation of NF-κB-mediated (proinflammatory) and type I IFN-mediated (antiviral) responses^73^. These two downstream pathways have distinct functions in the immune response^74^ and could potentially also differ in their roles on neurons. Since certain CpG-containing sequences activate only one of these pathways, while others activate both^75^, identifying the downstream TLR9 pathway(s) that mediate AAV-induced dendritic loss and changes in synaptic transmission could help identify specific CpG motifs that must be depleted. While straightforward CpG depletion was fully effective for protecting neurons from dendritic damage caused by AAV, these more targeted forms of CpG depletion and conversion may be useful in other contexts, either alone or in combination with other approaches to dampen TLR9 response, such as injecting lower viral titers^21,41,59^, AAV genome methylation^66,76^ or incorporation of exogenous TLR9 inhibitor sequences^71,72^.

In the current study, CpG-depleted AAV1*-pCAG-FLEX-EGFP* drove EGFP expression at levels comparable to those seen with unmodified virus. Similarly, a recent study that removed CpGs from the transgene region of an AAV8 designed to treat lipoprotein lipase deficiency found no change in therapeutic efficacy^77^, supporting the idea that AAV’s transgene region can be CpG depleted without compromising transgene function. Despite these and other examples of CpG depletion that did not reduce (or even enhanced^18,29^) transgene expression, it is possible that CpG depletion of some transgenes may disrupt transgene expression or function, even when synonymous mutations are made. Changes in mRNA stability, translational efficiency, and protein folding can allow synonymous mutations to impact protein expression and function in the absence of a change in protein sequence^51^. Thus, future studies using CpG depletion to prevent AAV-induced TLR9 responses will need to verify the expression and function of their transgene in their species and tissue.

Our *in vivo* validation of the protective value of CpG depletion against AAV-induced dendritic loss and synaptic weakening is limited by our use of only a single AAV vector (AAV1-*pCAG*-*FLEX*-*EGFP*). We cannot rule out the possibility that other AAV vectors, especially ones with higher CpG content, would require more extensive depletion of CpG motifs, including in regions outside of intronic and transgene regions. In this case, it may be useful to also consider depleting CpGs from promoter^18,29^ and/or ITR regions^58^, changes that have been made successfully without compromising the function of the encoded transgenes^18,29,58^. Additionally, the AAV genome used in this study is single-stranded (ssAAV), and self-complementary AAVs (scAAVs; recombinant AAV genomes that form double-stranded DNA) can induce a stronger TLR9-mediated innate immune response than single-stranded AAVs^78,79^. It will be important to determine if the CpG depletion approach used here can mitigate TLR9-mediated responses to scAAVs. Further refinement of the RF3 metric and the addition of tissue-specific codon optimization could improve the predictive power of CpG-AT in estimating the TLR9 activation potential of AAV genomes *in vivo*.

We created a web-facing application that provides tools to quantify, explore, and reduce CpG content (CpG-Assist Tool, or CpG-AT). The CpG content calculator allows users to quantify the CpG content of any ssDNA sequence and calculate its RF3 value. Users can also visualize the CpG content across the length of their sequence, making it easy to identify regions of high CpG density to target for replacement or depletion. The CpG depletion tool considers the codon usage of different species to create host-specific, codon-optimized, synonymous mutations in coding regions, and provides a resource to CpG-deplete intronic regions. The AAV component explorer aims to provide researchers with a tool to search and visualize the CpG content of a library of 148 AAV components, with the goal of making it easier to rapidly and affordably design AAVs with lower CpG content by selecting from readily available components.

Recent work has identified detrimental effects of AAV in the CNS, highlighting the need for strategies to protect neurons during AAV-mediated gene delivery. Here we find that depleting CpGs from the transgene and promoter fully prevents AAV-induced changes in circuit structure and function, without affecting transgene expression. We also find that the CpG content of commonly used AAV components varies widely, revealing that AAV’s CpG content can be minimized during virus design by selecting existing genome elements with lower CpG content. The CpG-AT application will allow users to quantify, explore, and minimize the CpG content of AAV genomes, facilitating the adoption of lower-CpG AAVs and increasing the safety and efficacy of AAV-mediated gene delivery.

## Materials and Methods

### 1. Animals

Male *C57BL/6J and Slc17a7-IRES2-Cre-D* (Strain # 037512) mice were obtained from Jackson Laboratory (The Jackson Laboratory, Bar Harbor, ME). All mice were acclimated for at least one week in the Princeton Neuroscience Institute vivarium before injections were performed. All AAV injections were performed on animals 8 to 10 weeks old, and tissues were harvested at 21 dpi (11-13 weeks old). Animals were group housed in Optimice cages (Animal Care Systems, Centennial, CO) with blended bedding (The Andersons, Maumee, OH) and enrichment (paper nesting and heat-dried virgin pulp cardboard hut). All mice were maintained on a 12hr light-dark cycle with ad libitum access to food (PicoLab Rodent Diet food pellets, LabDiet, St. Louis, MO) and water. All procedures were performed in accordance with protocols approved by the Princeton University Institutional Animal Care and Use Committee (IACUC) and in accordance with the animal welfare guidelines of the National Institutes of Health.

### 2. AAV Administration

Surgeries were performed as previously described^41,80^. Mice were anesthetized with isoflurane (5% for induction, followed by 1-2%, in 1 L/min oxygen; Viking Medical) and mounted into a stereotaxic instrument (David Kopf Instruments, Tujunga, CA). Body temperature was maintained and monitored using Kent Scientific PhysioSuite (Kent Scientific Corporation, Torrington, CT). An osmotic diuretic drug (15% D-mannitol in DPBS, 0.25 mL) was administered via intraperitoneal injection ten minutes prior to craniotomy.

Viruses were prepared using endotoxin-free plasmid prep kits, and virus preps were endotoxin-negative. Virus preps were treated with turbonuclease to minimize viral, host, and plasmid DNA impurities. 1×10^10^ vg of AAV2/1-*pCAG*-*FLEX*-*EGFP*-*WPRE* (serotype AAV1, using AAV2 ITRs; Princeton Neuroscience Institute, Addgene # 51502) were unilaterally injected into 3 individual brain regions (for a total of 3×10^10^ vg per brain) using borosilicate glass capillaries pulled on a Sutter Micropipette Puller (Model P-2000, Sutter Instrument Company). Injections of 67 nanoliters (nl) were made at 3 depths (500, 250, and 150 µm below the dura; 200 nl total) in regions corresponding to the motor (+2.46 to +1.42 mm from bregma), somatosensory cortex (+0.2 to -0.8 mm from Bregma and 2 mm lateral) and cerebellum (-5.40 to -8.24 mm from bregma) of the left hemisphere. All mice received a non-steroidal anti-inflammatory drug, ketoprofen, immediately post-surgery, 24 h, and 48 h post-surgery (0.2 mL, 50 mg/mL, subcutaneous). Untreated controls received no surgery, drug administration, or injection.

### 3. Diolistic Labeling and Sholl Analysis

Diolistic labeling and Sholl analyses to quantify pyramidal cell morphology were performed at 21 dpi as previously described^41^. Briefly, mice were anesthetized and intracardially perfused with 1.5% PFA diluted in PBS (Millipore Sigma Catalog # P3813-10PAK), and whole brains were removed. Brains were then post-fixed for one hour in 1.5% PFA and stored in cold PBS until used (usually 1-2 weeks). Brains were sectioned by vibratome (Leica Biosystems) at 100 μm and sections were stored in cold PBS. Tungsten particles, coated with the carbocyanine dye DiO, were delivered to cells in slices using the Helios Gene Gun System (BioRad) at 60-80 psi (pounds per square inch) of helium. Sections were kept at 4° C for 24 hours on a shaker and post-fixed in 4% formalin for one hour. Sections were counterstained with DAPI (1:1000 in PBS) for 5 minutes, washed, and coverslipped with ProLong Diamond Antifade Mountant (Invitrogen Catalog # P36961). Slides were imaged using a spinning disk confocal microscope (3I / Zeiss) and images gathered using Slidebook software (3I). Images were captured within 3 days of DiO delivery to minimize dye diffusion.

Traced cells were from layer IV-VI. Cells were selected by morphology and cortical layer; only cells with pyramidal-like somas in cortical layers IV-VI, a prominent apical neurite, and visible basal dendritic branches, within S1, were collected. In general, traced cells were the most complex cells from the cohort with the most complete and high quality DiO fill. The similarity of dendrites in untreated animals and in the contralateral hemisphere of AAV- and CpG-depleted AAV-injected animals suggests that consistent morphology can be obtained across animals, despite sampling a heterogeneous cell population and small differences in the plane of section for each brain.

After image collection, cells from all treatment groups were randomized into a cohort and labeled as “Cell 1”, “Cell 2” etc. before being transferred to a single folder. An investigator blind to treatment traced cells, quantified total dendritic length, and performed Sholl analyses on labeled neurons using the Simple Neurite Tracer Plugin in ImageJ Fiji. Dendritic length and Sholl intersections for cells in the contralateral hemisphere at 21 dpi were indistinguishable from untreated controls, indicating that AAV1-*pCAG*-*FLEX*-*EGFP* does not alter the morphology of pyramidal cells within the contralateral hemisphere at this age **(Supplemental Figure 1)**. Therefore, data are represented as a ratio of ipsilateral/contralateral within animal unless otherwise noted, to avoid influences from inter-animal variability in baseline brain structure, or minor differences in DiO intensity across bead batches.

### 4. Electrophysiology

#### Slice preparation

Acute coronal cortical slices were prepared from 11 to 14-week-old C57BL/6J male mice, either uninjected, or injected three weeks prior with unmodified or CpG-depleted AAV1-*pCAG*-*FLEX*-*EGFP*. Animals were anesthetized via inhalation isoflurane, and brains were rapidly dissected out. Slices (350 µm) were prepared using a Vibratome Series 1000 (Technical Products International) in ice-cold sucrose slicing solution, containing (in mM): 240 sucrose, 2.5 KCl, 10 Na-HEPES, 10 glucose, 1 CaCl, 4 MgCl, and 0.2 ascorbic acid, pH 7.3, bubbled with 100% oxygen. Slices were transferred to an incubation chamber containing 32° C high Mg^2+^ ACSF (in mM: 124 NaCl, 2.5 KCl, 26 NaHCO, 1.25 NaH PO, 10 Glucose, 1 CaCl, and 3 MgCl, equilibrated with 95% O_2_ / 5% CO_2_), and the temperature was allowed to gradually fall to room temperature (RT). Slices were incubated at RT for 1-3 hours before recording. For recordings, slices were transferred to an open recording chamber, submerged, and perfused at 2 mL/min with standard ACSF (in mM: 124 NaCl, 2.5 KCl, 26 NaHCO, 1.25 NaH PO, 25 Glucose, 2 CaCl, and 1.3 MgCl, equilibrated with 95% O_2_ / 5% CO_2_) at RT. Pharmacological agents were introduced by switching the perfusion reservoir.

#### Whole cell patch-clamp

Pipettes (4–8 MΩ) were filled with intracellular solution containing (in mM) 108 Cesium gluconate, 20 HEPES, 0.4 EGTA, 2.8 NaCl, 5 TEACl, 4 Mg-ATP, 0.3 Na-GTP, and 10 phosphocreatine, and 6.5 biocytin HCl, pH 7.3, 290 mOsm. Currents were detected using a Multiclamp 700 B amplifier (Axon Instruments), conditioned with a 2 kHz Bessel filter, and digitized at 5 or 10 kHz with a Digidata 1322A (Axon Instruments). Whole-cell input resistance and series resistance were monitored by hyperpolarizing voltage test pulses (− 5 mV) delivered before and after record acquisitions. Cells were excluded from analysis if the holding current (*I*_hold_) exceeded -100 pA, or if Rs was > 30 MΩ, or changed by >30% during the recording. mEPSCs were recorded from L4/5 pyramidal neurons of primary somatosensory cortex in the presence of 1 µm TTX (Hello Bio) and 100 µm Picrotoxin (Acros Organics). Each recording lasted a minimum of five minutes. Data was analyzed off-line using MiniAnalysis software (Synaptosoft, NJ, USA) and Origin 6.0 (Microcal Software, Inc.). The detection threshold for mEPSC was set at 6 pA. A total of 300 events were analyzed for each cell. For cumulative probability plots, at least 200 consecutive events were sorted.

### 5. Western blotting

Mice were anesthetized by 3% inhalation isoflurane and immediately decapitated for brain tissue extraction. Somatosensory cortex of ipsilateral hemispheres was isolated. Samples were homogenized in Pierce RIPA buffer (ThermoFisher Scientific catalog # 89901). Protein concentration was determined by bicinchoninic acid (BCA) analysis (ThermoFisher Scientific catalog # 23227). Prior to loading, proteins were diluted in 2x Laemmli sample buffer (Bio-Rad catalog # 1610737) containing a 1:20 dilution of 2-Mercaptoethanol (Bio-Rad catalog # 1610710) and boiled at 95°C for 5 minutes. 20 μg of protein was loaded into each lane of 12% Mini-PROTEAN TGX Precast protein gels (Bio-Rad catalog #4561043) and ran at 100V for 60 minutes in Novex Tris-Glycine SDS running buffer (diluted from 10x stock (catalog #LC2675)). Gels were transferred to .45 µm nitrocellulose membranes (Bio-Rad catalog # 1620145) at 350 mA for 60 minutes in Novex Tris-Glycine SDS running buffer containing 20% methanol. After transfer, membranes were washed in 1X TBS for 5 minutes, 3 times. Membranes were then blocked for 60 minutes at room temperature using a 5% milk (Biotium catalog # 22012) solution in TBS. After blocking, membranes were transferred to a primary antibody solution containing 1% milk in TBST (999 mL of TBS with 1 mL of Tween 20). Primary antibody dilutions were 1:1000 for anti-GFP antibody (Abcam catalog # ab290) and 1:1000 for anti-GAPDH antibody (Invitrogen, Fisher Scientific catalog # PIMA544398). Following overnight (16 hour) primary antibody incubation at 4°C, membranes were washed 3 times for 5 minutes with TBST and incubated in a secondary antibody solution containing 1% milk in TBST. LICOR IRDye secondary antibodies (catalog # 926-68071 & # 926-32210) were used at a 1:20,000 dilution. Following secondary antibody incubation for 60 minutes at room temperature, membranes were washed 5 times for 5 minutes with TBS. Membranes were imaged using a LICOR Odyssey Fc Imager.

### 6. CpG depletion

CpG dinucleotides were depleted from the transgene and intronic regions of AAV1-*pCAG*-*FLEX*-*EGFP*-*WPRE*. In the transgene coding region, we replaced CpG dinucleotides by creating synonymous, species-specific, codon-optimized mutations. To determine which codon(s) would replace CpG dinucleotide-containing (example: CGG) or CpG dinucleotide-spanning (CUC-GGG) sequences, we used the updated codon usage tables found in the High-performance Integrated Virtual Environment-Codon Usage Tables (HIVE-CUTs) database^47^. We replaced the current CpG -containing or -spanning dinucleotide with codon(s) that did not change the encoded amino acid(s) and had the highest usage in mice. If the highest usage codon was being replaced, we replaced it with the next-highest usage codon. In intronic regions of AAV1-*pCAG*-*FLEX*-*EGFP*, we removed CpG dinucleotides by replacing the C with an A. We have made this species-specific CpG depletion pipeline available via our online application (see CpG-AT application). The pipeline available on CpG-AT has one modification to the CpG depletion protocol used here: in intronic regions, CpG-AT will change the C in CG motifs to a G if the preceding base is not a C, to keep GC content as consistent as possible, which may help with viral production and/or transgene expression^51,58^. In cases where the preceding base is a C, the C from the CG motif is changed to an A.

### 7. Statistics

Statistical analyses were performed using GraphPad Prism. For all datasets, a Shapiro-Wilk test was performed to ensure data were normally distributed. A two-tailed Grubb’s test was used to identify potential outliers. For all cases where 2 or more comparisons were made, a Brown-Forsythe test for homogeneity of variance was conducted to test for the assumption of equal variances. If data met the equal variance assumption (all figures except **Figure 3B**), a standard one-way ANOVA was conducted followed by Tukey’s multiple comparison test. If data did not meet the equal variance assumption (significant result in the Brown-Forsythe test; **Figure 3B**), a Brown-Forsythe one-way ANOVA test was conducted followed by Dunnett’s T3 multiple comparison test.

### 8. Application

The CpG-AT application was created using the Shiny package for Python and is available at https://cpgat.princeton.edu/.

## Data Availability Statement

The datasets generated and analyzed during the current study are available from the corresponding author upon request.

## Acknowledgements

We wish to thank Oliver Huang, Angela Chan, and the Princeton Viral Vector Core for their technical assistance during these experiments. Research reported in this publication was supported by the National Institute of General Medical Sciences of the National Institutes of Health under grant number T32GM007388 (C.D.), the New Jersey Commission on Brain Injury Research under grant number CBIR25FEL014 (C.D.), and the Innovation Fund for New Ideas in the Natural Sciences (L.M.B.). The content is solely the responsibility of the authors and does not necessarily represent the official views of the National Institutes of Health. Some images created in BioRender.

## Author Contributions

C.D. and L.B. conceived the study and wrote the paper. C.D. designed and created the CpG-depleted virus. C.J. performed AAV injections and B.C. and C.D. collected tissue for analysis.

C.D. designed figures and performed data analysis. C.D. and B.C. performed DiOlistic labeling and confocal fluorescence imaging. C.D. and B.C. performed blinded dendritic length and Sholl analyses. C.D. conducted western blotting procedure. J.M. performed electrophysiology experiments. A.V.N and C.D. performed calculations of CG content. C.D. designed and created the CpG-AT application. C.D. and G. C-H. tested and validated the CpG-AT application.

