## Supplementary Material for "Adeno-Associated Virus-Induced Neurotoxicity is Prevented by CpG Depletion"

**Supplementary Table 1: Library profiling the CpG content of 148 AAV components.**

The total CpG dinucleotides, % CpG dinucleotides, and RF3 values of AAV components are provided here, along with corresponding source and reference information. AAV components are grouped by category (transgene, promoter/enhancer, miscellaneous, and ITR) and sub-category (biosensor, chemogenetics, fluorescence, optogenetics, other (transgene), recombinase, broad promoter, specific promoter, inducible promoter, histone, linker, regulatory element, and tag).

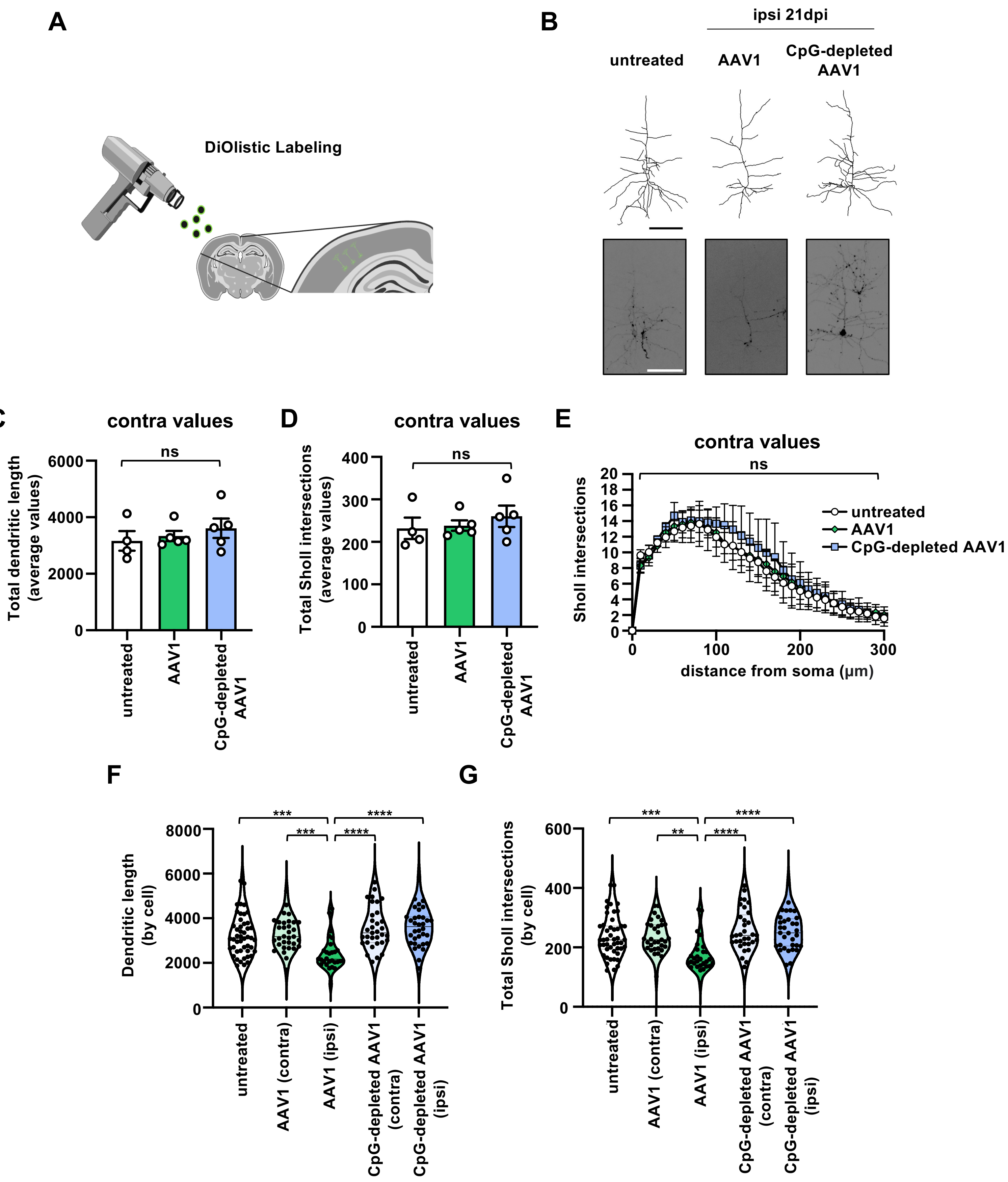
A
DiOlistic Labeling
B
ipsi 21dpi
untreated
AAV1
CpG-depleted AAV1
C
contra values
ns
Total dendritic length (average values)
6000
4000
2000
0
untreated
AAV1
CpG-depleted AAV1
D
contra values
ns
Total Sholl intersections (average values)
400
300
200
100
0
untreated
AAV1
CpG-depleted AAV1
E
contra values
ns
Sholl intersections
untreated
AAV1
CpG-depleted AAV1
distance from soma (μm)
0
100
200
300
F
Dendritic length (by cell)
***
***
****
****
8000
6000
4000
2000
0
untreated
AAV1 (contra)
AAV1 (ipsi)
CpG-depleted AAV1 (contra)
CpG-depleted AAV1 (ipsi)
G
Total Sholl intersections (by cell)
***
**
****
****
600
400
200
0
untreated
AAV1 (contra)
AAV1 (ipsi)
CpG-depleted AAV1 (contra)
CpG-depleted AAV1 (ipsi)

**Supplementary Figure 1. Methods for quantification of dendritic complexity, and raw values for uninjected contralateral hemispheres.**

**(A)** Schematic demonstrating the DiOlistic labeling procedure used to quantify dendritic complexity. DiO-coated bullets are ballistically fired into brain sections thereby sparsely labeling neurons with the fluorescent DiO dye (see Methods for details). **(B)** Examples of DiO-filled cortical neurons (bottom) and corresponding tracings (top) from untreated wild type mice or ipsilateral (injected) hemispheres of wild type mice injected with $1x10^{10}$ vg of either unmodified or CpG-depleted AAV1-*pCAG-FLEX-EGFP*. DiO-containing images shown are max intensity z-projections of z-stacks composed of 50 images separated by 2 µm each. Note: tracing was performed by moving through the z-stack (not using the max intensity z-projection). Scale bar = 100 µm. **(C)** Average total dendritic length and **(D)** total Sholl intersection values in C57BL/6J untreated control mice or contralateral (uninjected) hemispheres from C57BL/6J mice injected with $1x10^{10}$ vg of either unmodified AAV1-*pCAG-FLEX-EGFP* or CpG-depleted AAV1-*pCAG-FLEX-EGFP* (same mice from **Figure 2**). **(E)** Contralateral Sholl intersections over increasing distance from the soma for cortical neurons from the same animals as in C-D**. (F-G)** Raw total dendritic length and total Sholl intersection values for individual cells from unmodified AAV1-*pCAG-FLEX-EGFP*- and CpG-depleted AAV1-*pCAG-FLEX-EGFP*-injected animals from the same animals as in C-D**.** *n=4* untreated control mice, 10-14 neurons per animal (average = 12 cells)*; n*=5 mice injected with unmodified AAV1-*pCAG-FLEX-EGFP,* 10-14 neurons per animal (average = 12 cells); *n*=5 for mice injected with CpG-depleted AAV1-*pCAG-FLEX-EGFP,* 12-15 neurons per animal (average = 13 cells). Data in B-E represented as mean ± SD. ****$p < .0001$; ***$p < .001$; **$p < .01$; ns, not significant. For **(F-G)**, all statistical comparisons not specified are not significant. Statistical analysis in **(C D;F-G)**: one-way ANOVA with Tukey's multiple comparison test, in **(E)**: two-tailed unpaired t-test.

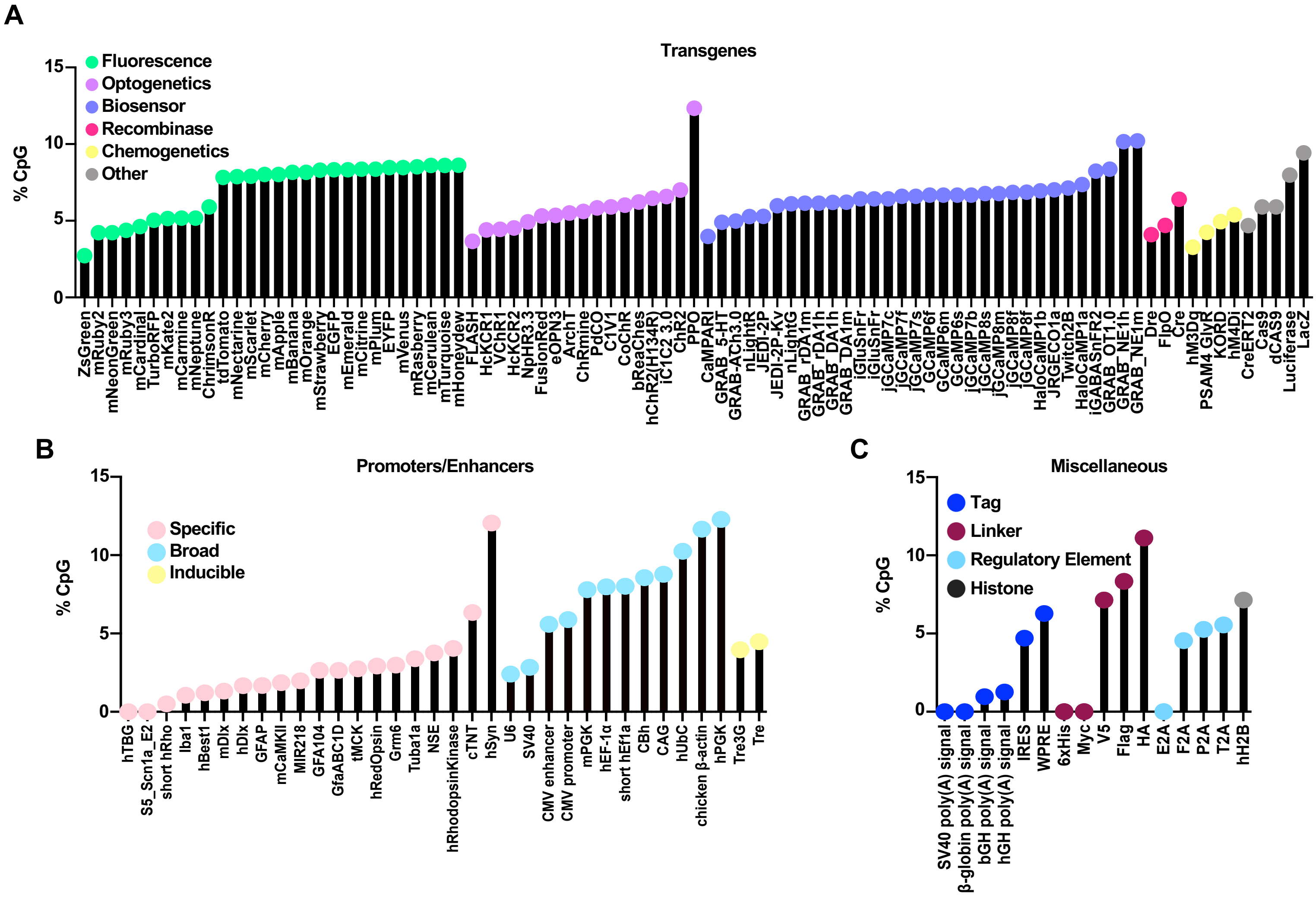


**Supplementary Figure 2. % CpG dinucleotides of commonly used AAV transgenes, promoters, enhancers, and miscellaneous components.**

% CpG dinucleotides of **(A)** transgenes classified by general function: fluorescence, optogenetics, biosensor, recombinase, chemogenetics, other; **(B)** promoters/enhancers classified by broad, specific, and inducible; and **(C)** miscellaneous components classified by tags, linkers, regulatory elements, and histones.

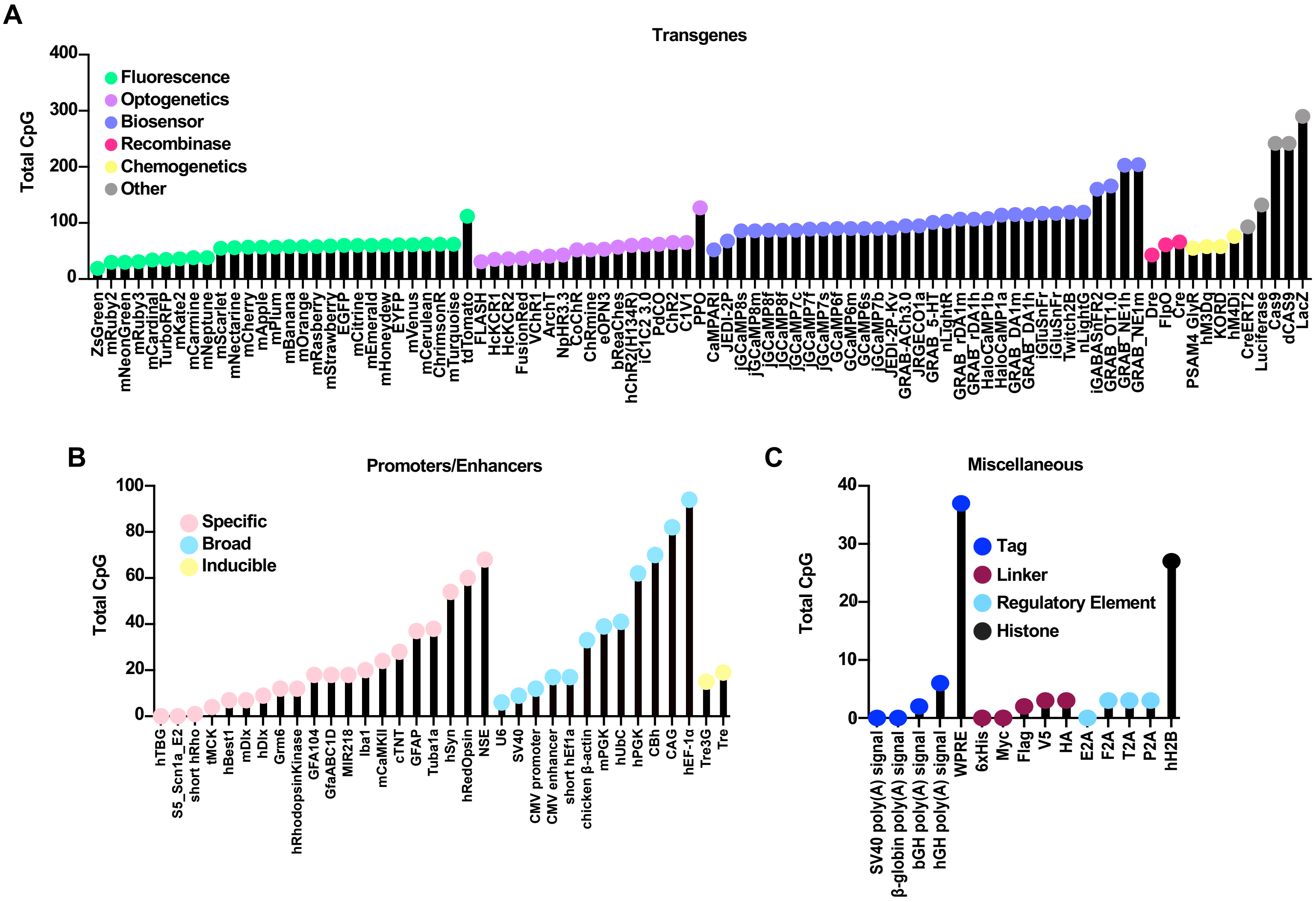


**Supplementary Figure 3. Total CpG dinucleotide count of commonly used AAV transgenes, promoters, enhancers, and miscellaneous components.** Total CpG dinucleotide count of **(A)** transgenes classified by general function: fluorescence, optogenetics, biosensor, recombinase, chemogenetics, and other; **(B)** promoters/enhancers classified by broad, specific, and inducible; and **(C)** miscellaneous components classified by tags, linkers, regulatory elements, and histones.

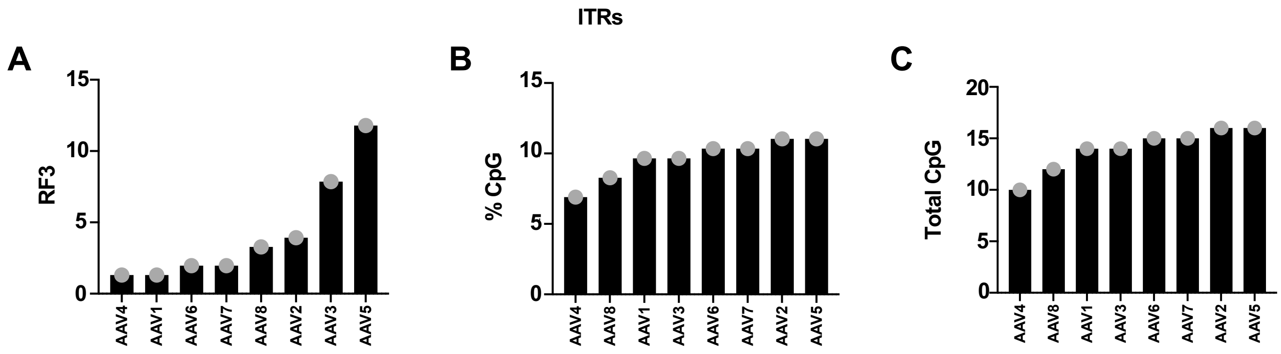


**Supplementary Figure 4. RF3, % CpG, and total CpG dinucleotide count of ITRs. (A)** RF3, **(B)** % CpG, and **(C)** total CpG dinucleotide count of ITRs (bases 1-145 of full AAV genome sequences).
